# A Tunable, Simplified Model for Biological Latch Mediated Spring Actuated Systems

**DOI:** 10.1101/2020.12.02.408740

**Authors:** Andrés Cook, Kaanthi Pandhigunta, Mason A. Acevedo, Adam Walker, Rosalie L. Didcock, Jackson T. Castro, Declan O’Neill, Raghav Acharya, M. Saad Bhamla, Philip S. L. Anderson, Mark Ilton

**Affiliations:** Department of Physics, Harvey Mudd College, Claremont, CA 91711, USA; School of Chemical and Biomolecular Engineering, Georgia Institute of Technology, Atlanta, Georgia 30318, USA; Department of Evolution, Ecology, and Behavior, University of Illinois at Urbana-Champaign, Urbana, IL 61801, USA

## Abstract

We develop a model of latch-mediated spring actuated (LaMSA) systems relevant to comparative biomechanics and bioinspired design. The model contains five components: two motors (muscles), a spring, a latch, and a load mass. One motor loads the spring to store elastic energy and the second motor subsequently removes the latch, which releases the spring and causes movement of the load mass. We develop open-source software to accompany the model, which provides an extensible framework for simulating LaMSA systems. Output from the simulation includes information from the loading and release phases of motion, which can be used to calculate kinematic performance metrics that are important for biomechanical function. In parallel, we simulate a comparable, directly actuated system that uses the same motor and mass combinations as the LaMSA simulations. By rapidly iterating through biologically relevant input parameters to the model, simulated kinematic performance differences between LaMSA and directly actuated systems can be used to explore the evolutionary dynamics of biological LaMSA systems and uncover design principles for bioinspired LaMSA systems. As proof of principle of this concept, we compare a LaMSA simulation to a directly actuated simulation that includes a either Hill-type force-velocity trade-off or muscle activation dynamics, or both. For the biologically-relevant range of parameters explored, we find that the muscle force-velocity trade-off and muscle activation have similar effects on directly actuated performance. Including both of these dynamic muscle properties increases the accelerated mass range where a LaMSA system outperforms a directly actuated one.

## INTRODUCTION

A diverse array of organisms use stored elastic energy to drive rapid movements. These organisms use motors, springs, and latches to perform a latch mediated spring actuated (LaMSA) motion, and remarkably, they can use this mechanism to outperform current engineering design for repeatable motion at small size-scales [1]. Models have been developed to understand the extreme biomechanics of latch-mediated spring actuated organisms. Organism-specific models, including both continuum mechanics-based models [2–10] and physical modeling with biomimetic devices [2, 10–14], have been used to test hypotheses about the movement of specific organisms (Table I summarizes examples of recent work).

**TABLE I.**
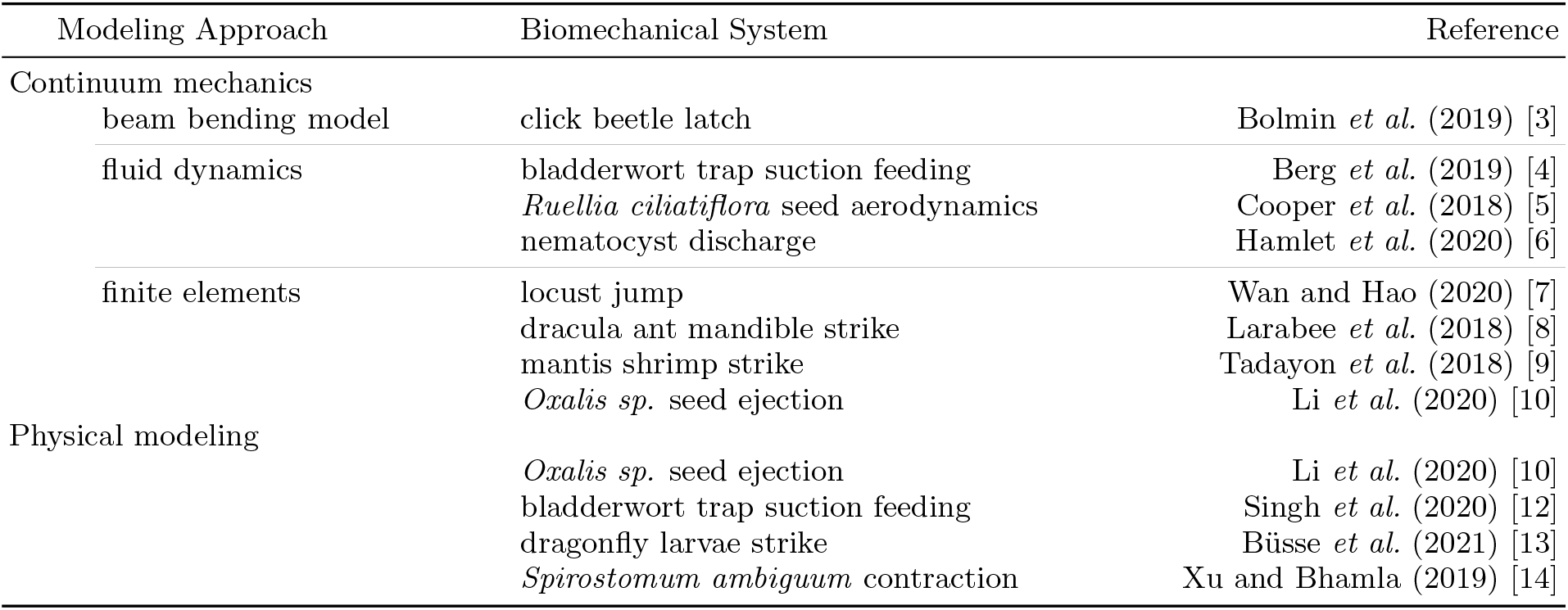
Recent examples (since 2018) of modeling latch-mediated spring actuated organisms, which includes both mathematical and physical approaches. For a review of earlier work see ref. [27].

In contrast to organism-specific models, ‘simple models’ with reduced complexity [15] are primarily used for making inter-species comparisons, and for testing scaling relationships and the sensitivity of kinematic performance to different characteristics of the organism. These simple models can also have broad applicability and enable the rapid testing of ideas [15], and typically include muscle motors, springs, masses, and other mechanical linkages. In recent work, these models have been applied to jumping organisms [16–23] and augmented human movements [24, 25]. General models have also been used to test hypotheses about the scaling and effectiveness of biological spring mechanisms [23, 26–29]. These types of models have similarities to template models – simple biomechanical models that demonstrate a particular mechanical behavior [30].

In our previous work [27], we used a simplified mathematical model to illustrate trade-offs between the components of a general LaMSA system. The components of a LaMSA system (the latch, spring, loading motor, and load mass) were modeled as a simplified mechanical system and given material, geometric, and dynamic properties; however, the properties of the system components were limited to motors and springs with linear properties, specific latch shapes, frictionless interactions between components, and a fixed unlatching velocity.

Here we develop a LaMSA Template Model with accompanying open-source software. Our model here includes a more general framework for defining LaMSA components (refs. [23, 26, 27, 29] are all particular cases of this new model). This broader approach allows the model to be tuned to a specific organism, group of organisms, or a biological scaling relationship to explore questions in comparative biomechanics and LaMSA system design. Our approach also includes non-linear and time-dependent material properties. Additionally, we provide a generalized treatment of the latch that includes friction, allows for different latch shapes, and includes an unlatching motor that drives the latch removal of the system, similar to the one recently hypothesized to occur in some biological systems [13].

Finally, as an example of this LaMSA model’s utility, we use the model to explore how dynamic muscle properties affect the power output of both a LaMSA system and a system where the muscle is used to directly actuate movement. Two important dynamic aspects of muscle are a force-velocity trade-off (the muscle exerts less force at higher velocities) and an activation rate (it takes some time for the muscle to reach its maximum force) [31]. Previous work has been focused on how muscle force-velocity trade-offs limit power output for a directly actuated system [26, 27]. This force-velocity trade-off is a principal reason LaMSA systems can outperform comparable muscle-driven ones at small load mass; however, it is unclear how significant this force-velocity effect is compared to the activation dynamics of muscle. Here we directly compare the effect of the muscle force-velocity trade-off to the effect of muscle activation. Using the LaMSA Template Model with inputs guided by biologically-relevant sizes and masses, we find that the muscle force-velocity trade-off and activation dynamics cause a similar reduction in directly actuated kinematics. Combining the two effects, the mass range where a LaMSA system outperforms a directly actuated one increases by a factor of *≈* 5 times compared to systems where only one of the two time dependent motor properties is included.

## METHODS

### LaMSA Template Model

In our model, the motion of a LaMSA system is comprised of three distinct phases: loading, unlatching, and spring actuation. In the loading phase (Fig. 1A, first panel), a loading motor (e.g. muscle) deforms a spring starting from the spring’s stress-free equilibrium length. We make the simplifying assumption that the loading occurs slowly enough to approximate it as a quasi-static motor contraction – i.e. the loading follows the isometric force-length curve in the case of a muscle motor. The loading phase down (in the *− y* direction) matches the spring force pushing up. After the loading phase, the loading motor is removed from the system and the spring is held in place by a latch (Fig. 1A, second panel). The second phase of motion, the unlatching phase (Fig. 1A, third panel), begins with the activation of an unlatching motor that pulls the latch out of the way. During the unlatching phase the load mass and latch undergo a complex interaction. The interaction between the load mass and latch is modeled as a frictional contact between two rigid bodies, and the unlatching phase ends when there is no longer any contact between the load mass and latch. Once the contact breaks, the load mass is actuated solely by the spring, which undergoes a rapid expansion (Fig. 1A, fourth panel). Spring actuation continues until the spring returns to its equilibrium length where it no longer applies a force to the load mass, which causes the load mass to undergo take-off (Fig. 1A, fifth panel). In the model, we assume that the latch shape is sufficiently smooth that after the latch disengages, it does not re-engage at a later time. This assumption enables the clear delineation of the unlatching and spring actuation phases.

**FIG. 1.**
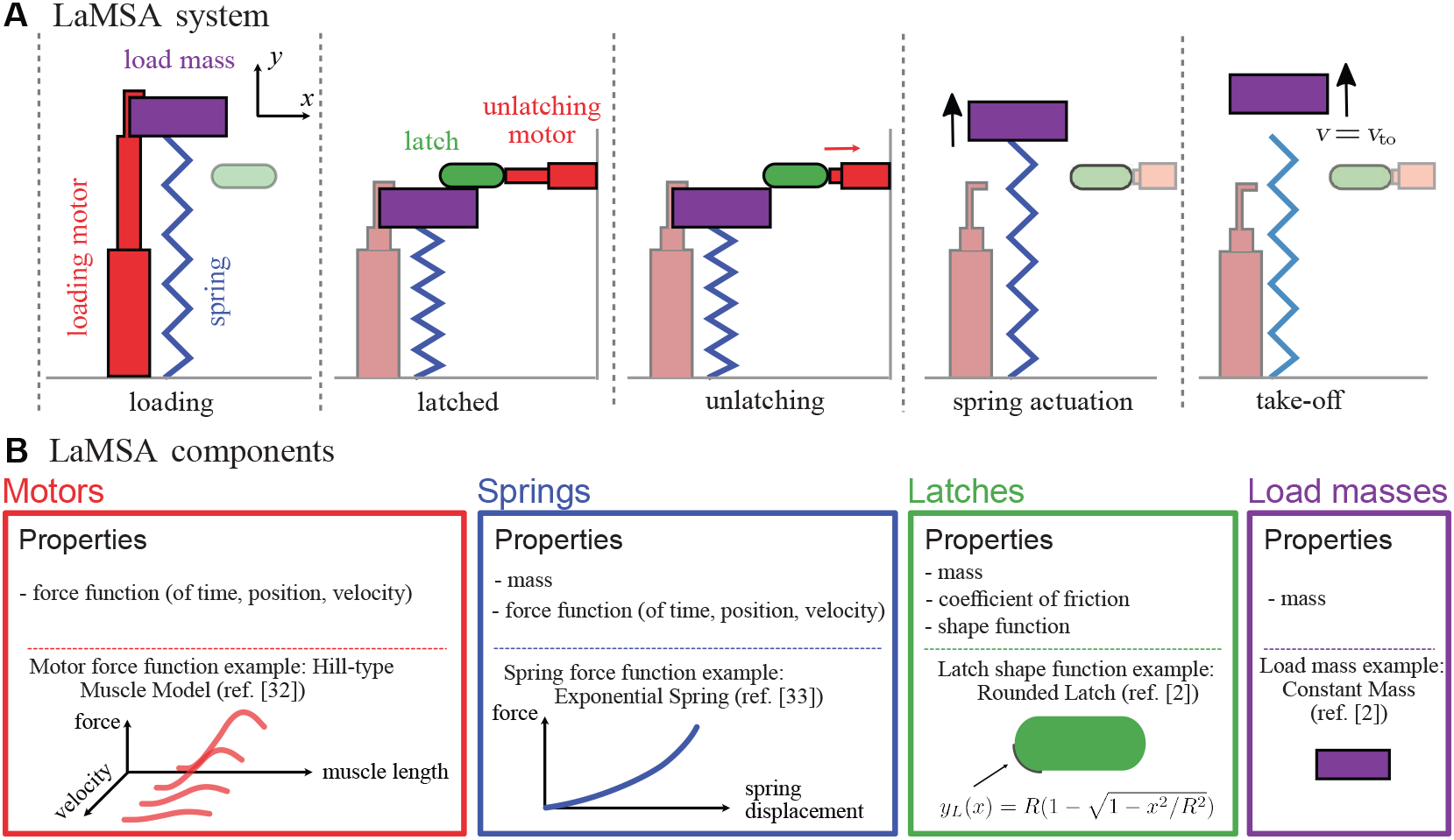
Schematic description of the LaMSA Template Model with a loading motor, spring, latch, unlatching motor, and load mass. **A** The sequence of important events during the movement of a LaMSA system, which includes three delineated phases of motion in the model: loading, unlatching, and spring actuation (diagram modified from ref. [27]). **B** The properties of the components used in the LaMSA Template Model, and an example of each component that is explored in this work (see Table A1 in Appendix A for the specific functions and parameters used in this manuscript).

The dynamics of a LaMSA system depends on its components and the interaction between them. In our model, these components are classified into motors, springs, latches, and load masses (Fig. 1B). Each component is constrained to move along a single coordinate axis in the model (the loading motor, load mass and spring move along the *y* axis; the latch and unlatching motor move along the *x* axis). We develop our model with the aim to give general properties to each component. The motors and springs in the LaMSA system are characterized by their force output. The loading motor force (*F*_lm_), the unlatching motor force (*F*_um_), and spring force (*F*_sp_) are all assumed to be functions of time, displacement, and velocity. Latches are given a shape function *y*_*L*_(*x*) that describes the geometry of the latch. The shape function relates horizontal motion of the latch (in the *x* direction) to vertical displacements of the load (in the *y* direction). For example, the rounded latch used in this work, which has circular edges of radius *R*, has a shape function shown in Fig. 1B. The shape function describes the shape of the latch where it contacts the load mass. The derivatives of this shape function with respect to *x* determine the latch slope function 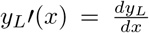 and latch concavity 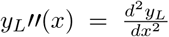. The functions describing shapes and forces are taken as inputs into the model to allow for hypothesis testing of non-linear properties. In addition, the mass of the system can be distributed in the spring mass (*m*_*s*_), latch mass (*m*_*L*_), and load mass (*m*). With these definitions, we lay out the mathematical description of the model according to its three phases of motion.

#### LaMSA Template Model: Loading Phase

In the loading phase, the loading motor slowly applies a force causing a displacement of the spring. The final displacement of the spring at the end of the loading phase, *y*_0_, is the displacement in which the loading motor force and spring force are equal and opposite, namely

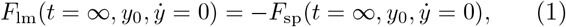

where 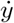 is the velocity in the *y* direction. The condition that *t* = *∞* and 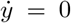 corresponds to a slow, quasistatic loading of the spring. The loaded displacement, *y*_0_, depends on how the force-displacement properties of the loading motor and spring interact.

#### LaMSA Template Model: Unlatching Phase

The unlatching phase starts with the activation of the unlatching motor at time *t* = 0. The load mass starts with an initial position *y* = *y*_0_ and velocity 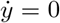, while the latch has an initial horizontal position *x* = 0 and velocity 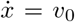. By analyzing the spring force pushing on the load mass, the unlatching motor force pulling on the latch, and the contact force between the load mass and latch, we derive that the differential equation for the acceleration of the latch, 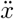, during the unlatching phase of motion

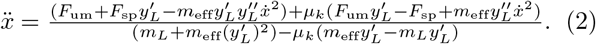

The term *m*_eff_ in Eq. (2) is the effective mass for the system, with *m*_eff_ = *m* + *m*_*s*_*/*3 [27], and *μ*_*k*_ is the coefficient of friction between the latch and load mass. A full derivation of Eq. (2) is presented in Appendix B. For a LaMSA system undergoing rotational motion, the effective mass and mapping onto Eq. (2) is provided in Appendix C. From the dynamics of the latch and the latch shape, the acceleration of the load mass during the unlatching phase is given by taking a chain rule, namely

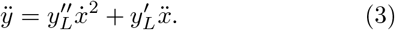

To determine the end of the unlatching phase, we solve for the magnitude of the normal component of the contact force between the load mass and latch,

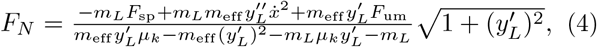

and require that this magnitude be *F*_*N*_ *≥* 0 during the unlatching phase to ensure there is still contact between the load mass and latch. Therefore, we solve for when *F*_*N*_ = 0 to determine the unlatching duration *t*_*L*_, which marks the end of the unlatching phase and the beginning of the spring actuation phase of motion.

#### LaMSA Template Model: Spring Actuation Phase

After unlatching, the load mass undergoes a purely spring-driven motion given by

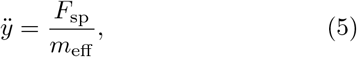

where the spring force can depend on position, velocity, and time. The initial conditions for this phase are given by the ending condition from the unlatching phase. Namely, for the spring actuation phase, the initial position of the load mass is *y*(*t* = *t*_*L*_) and its initial velocity is 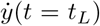. The spring actuation phase ends when the spring stops pushing on the load mass - i.e. when *F*_sp_ = 0.

### Direct Actuation Model

The direct actuation model uses the loading motor of the LaMSA system to directly drive the load mass. To ensure the motor in the directly actuated model is being used in a comparable way to the LaMSA model, the mass is accelerated by the motor using a motor contraction. Therefore, the equation of motion for the load mass is given directly by the force applied by the motor as it contacts,

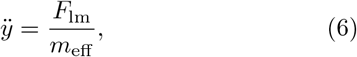

where the loading motor force can depend on position, velocity and time. The initial condition for the directly actuated system is that the motor and load mass are initially at rest, with the motor at its undisplaced initial length. Take-off occurs when the load mass reaches its maximum velocity and *F*_lm_ = 0.

### LaMSA and Direct Actuation Software Implementation

The LaMSA and direct actuation models were implemented in MATLAB. This software implementation is open-source and available at https://posmlab.github.io [32]. The software allows a user to select a LaMSA system from a library of components (motors, springs, latches, and load masses), set parameters for each component, and run a simulation to determine the dynamics of that system (as both a LaMSA system and a directly actuated system). The software can be used to iterate over the LaMSA system component parameters (e.g. spring stifness) and rapidly generate the dynamics for variety of LaMSA systems.

### Model Input Parameters

The inputs parameters to the model were chosen based on the accelerated mass, characteristic velocities, and typical accelerations of the larger biological LaMSA listed in the supplementary materials of ref. [27]. To explore the role of the dynamic properties of muscle, we used a Hill-type muscle motor based on ref. [31], which is one of the default components included in the LaMSA Template Model software. A muscle activation rate of 200 s^*−*1^ was chosen as a typical rate based on the force generation delay of small animals reported in ref. [33]. The full list of parameters used in this work are reported in Table A1.

## RESULTS AND DISCUSSION

Using the components and parameters in Table A1, the output from a single simulation generated using the software is shown in Fig. 2. The software output includes information about the loading phase, and the dynamics of the latch and load mass during the unlatching and spring actuation phases. For the load mass dynamics, the simulation generates the position *y*(*t*), velocity 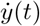, and forces acting on the load mass. From the position and velocity of the load mass, commonly used metrics for kinematic performance in biomechanics (e.g. maximum acceleration and maximum power [1]) are calculated. The maximum load mass acceleration (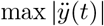, calculated from the numerical derivative of 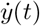 and maximum power delivered to the load mass 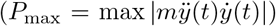 depend on the input parameters to the model, and the open-source software enables a rapid iteration over a range of input parameters.

**FIG. 2.**
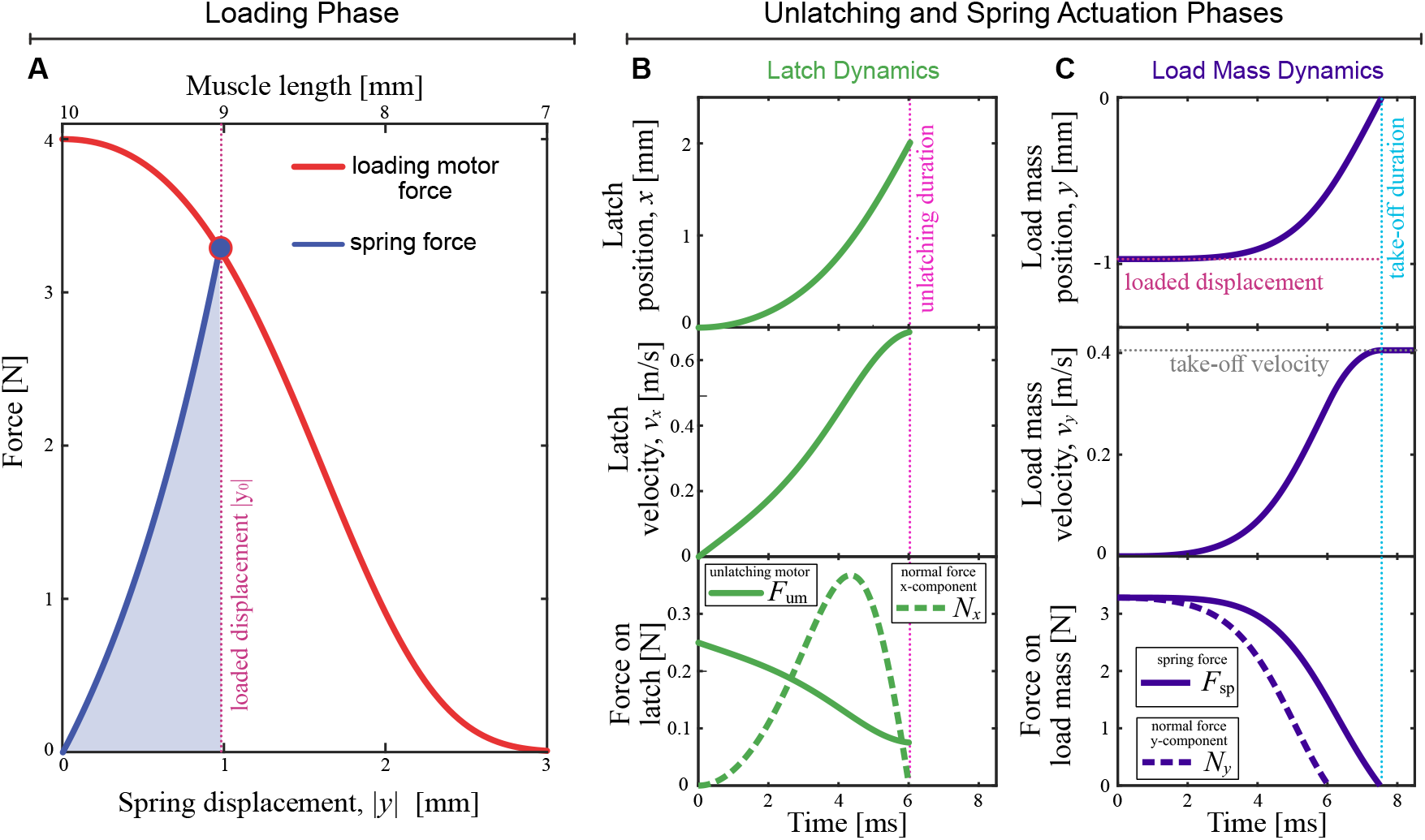
Example output from the model using the components and the biological LaMSA parameters listed in Table A1. **A** The force-length curve for a Hill-type muscle motor loading a tendon-like exponential spring. The LaMSA system loads up to a spring displacement *y*_max_ calculated by equating the loading motor and spring forces. **B-C** The dynamics during the unlatching and spring actuation phases for the latch (panel B) and load mass (panel C). The end of the unlatching phase is marked by the pink vertical dotted line showing the unlatching duration (*t*_*L*_ *≈* 6 ms), which occurs when the normal force *N* between the latch and load mass goes to zero (dashed curves in B-C). After unlatching, the load mass is actuated solely by the spring up until take-off duration (*t*_to_ *≈* 7.5 ms) when the spring force goes to zero, and the load mass reaches its take-off velocity (*v*_to_ *≈* 0.4 m*/*s).

For a motor directly actuating a load mass, the maximum power output depends on accelerated mass, with an upper bound set by the dynamic properties of the motor (Fig. 3, red curves). Driving the mass with a motor that has only a force-velocity trade-off (setting *r*_act_ = *∞* and *v*_max_ = 5 m*/*s in the model) has a similar effect to a motor that only has activation dynamics (setting *r*_act_ = 200 s^*−*1^ and *v*_max_ = *∞* in the model). Both motors reach an upper bound on their maximum power output when driving small masses (Fig. 3, dashed and dotted red curves). Therefore, even in the absence of a force-velocity trade-off, motors with slow activation rates still have performance limitations when driving small masses. Including both the effects of force-velocity and activation in the motor, as projectile mass is decreased the maximum power output of a directly actuated movement not only saturates to a maximum value, but further decreases for the smallest masses (Fig. 3, solid red curve).

**FIG. 3.**
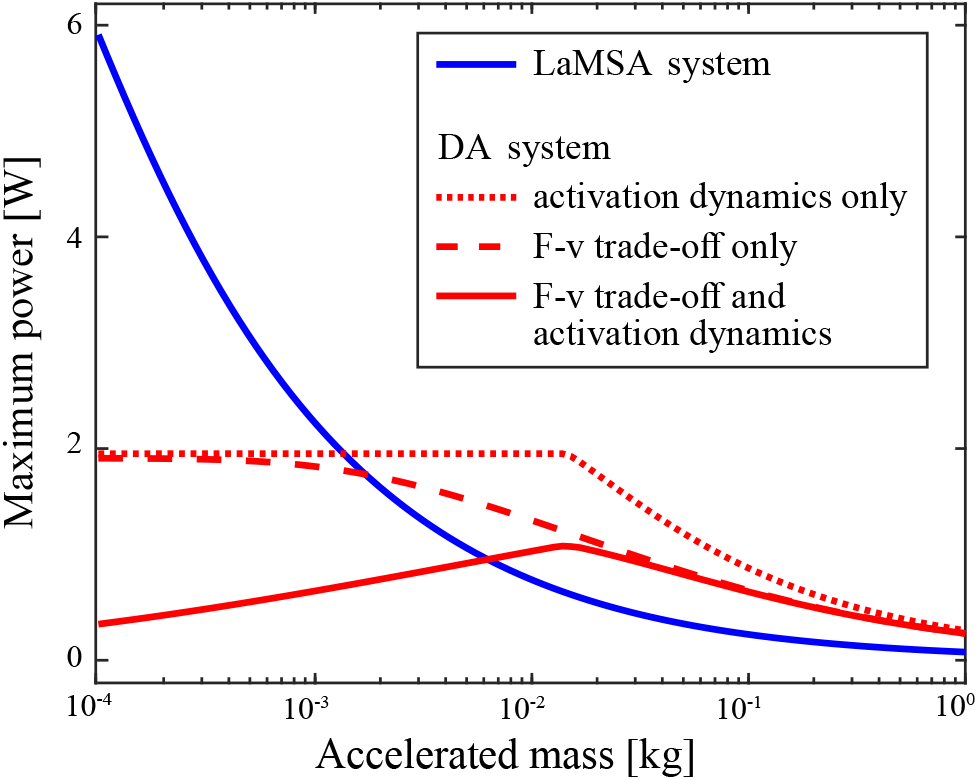
Both a motor’s activation rate and its force-velocity trade-off affect its maximum power output when it directly actuates a projectile. Compared to a using a motor in a LaMSA system (blue solid curve), the maximum power output of a directly actuated system (red curves) is worse for smaller masses. A motor that has both a force-velocity and activation limitation (solid red curve) has a significantly reduced performance at low masses compared to one with only a force-velocity trade-off (dashed red curve) or only an activation rate limitation (dotted red curve). The intersection between the LaMSA and directly curve shifts to a higher mass when both dynamic effects of the motor are included.

In contrast to the directly actuated systems, the LaMSA system is insensitive to the force-velocity trade-offs and activation dynamics of the loading motor. Varying the loading motor in the LaMSA system using the same three conditions as the directly actuated one (activation dynamics only, F-v trade-off only, F-v trade-off and activation dynamics), the maximum output for those three LaMSA systems is identical (Fig. 3, solid blue curve). The independence of the LaMSA system on the dynamic properties of the loading motor is a result of the slow, quasi-static loading assumption made in the model. This assumption is justified for biological LaMSA systems like mantis shrimp where typical loading rates are orders of magnitude slower than the rate of elastic energy release [34], but the loading motor dynamic properties can be important when considering simultaneous loading and release of a series elastic system [26].

The power output of comparable LaMSA and directly actuated systems have a mass dependent transition that is affected by the dynamic properties of the motor. Comparing the three different motor conditions in Fig. 3, the crossovers between the power output of the directly actuated and LaMSA systems is shifted to a larger mass (by a factor of *≈* 5 times) when both the force-velocity and activation dynamics of the motor are included in the simulation. This result suggests that in systems where there is a development and transition of a LaMSA mechanism (e.g. in some species of mantis shrimp [35]), care should be given to both muscle force-velocity and activation dynamics when modeling the transition from LaMSA to directly actuated movement.

Beyond this proof of principle example, the LaMSA Template Model and open-source software provides an extensible platform for exploring biological LaMSA systems. Although this model was formulated generally to encompass a broad range of LaMSA systems, the model can be tuned to specific biological systems because of the flexibility in how system components are defined. The relevant range of input parameters and any interdependence between them can be informed by observed biological data and scaling. For example, depending on the system, the characteristic lengths of the system (i.e. muscle lengths, latch radius, spring length) could be constrained in the model to follow an isometric scaling. These constraints on the model can be used make inter-species comparisons and to investigate to what extent kinematic performance changes over the course of development for a given species.

Flexible component definitions also enables new components to be created that address specific biological questions. For example, Deban *et al*. performed a comparative analysis of tongue projection across salamander species which actuate their tongue projection with a LaMSA mechanism or by direct muscle actuation [36]. The LaMSA projection mechanism can not only lead to higher kinematic performance, but is also robust to temperature variations [36]. To explore this system with the LaMSA model presented here, new components can be created in the software that introduce a temperature-dependent motor and spring. Adding these components would yield a theoretical prediction of the relative sensitivity of the tongue projection performance to temperature for the two groups of salamanders. Comparing this prediction to the observed kinematics could be used to inform the modeling of how biological motors and springs depend on temperature.

Finally, the model and software presented here can offer insights into how the interrelationships between input parameters and performance may influence the evolution of these biological systems via the concept of mechanical sensitivity. Mechanical sensitivity refers to the idea that variation between parts of a multi-part system are not necessarily equal in relation to their influence on the output of the system [37, 38]. Applied to a LaMSA system, we might hypothesize that variation in the spring would result in a larger variation in maximum power than variation in the latch mass. If so, that could mean that the latch mass has more freedom to evolve without altering performance. Such patterns have been identified in both mantis shrimp and fish [38, 39] and have been shown to influence rates of morphological evolution [40–42]. The model presented here offers an opportunity to quantitatively map how shifts in input parameters affect multiple performance metrics simultaneously, allowing for a comprehensive analysis of mechanical sensitivity.

## CONCLUSION

The LaMSA Template Model and open-source software presented here balances modeling principles of simplicity and extensibility. Simplicity is provided by making explicit assumptions about how the components are connected in the model, and extensibility is achieved though flexibility of defining the individual components. With these principles, the model enables the rapid testing of ideas by simulating kinematic output across the varying model parameters. This model also opens possible new directions for future work by providing a framework for others to build upon. Case studies using the model will inform best practices for tuning the model to explore a specific biological system. Exploring biological and bioinspired LaMSA systems with this model will require input from members of comparative biomechanics community through the use of the open-software (available at https://posmlab.github.io [32]), requesting new features, and actively contributing to software development.

## AUTHOR CONTRIBUTIONS

AC and MI designed the research; all authors contributed to the model development; AC, KP, MAA, AW, RLD, JTC, DO, RA, and MI wrote the software; AC and MI wrote a first draft of the manuscript; all authors revised and edited the manuscript.

## ACKNOWLEDGMENTS

This material is based upon work supported by the National Science Foundation under Grant No. 2019371. We thank the Harvey Mudd College Physics Summer Research Fund and the N. Sprague III Experiential Learning Fund for financial support. The authors thank S.N. Patek and Justin Jorge for stimulating discussions and helpful suggestions on this work.

## APPENDICES

### Appendix A: Table of Parameters Used

**TABLE A1.**
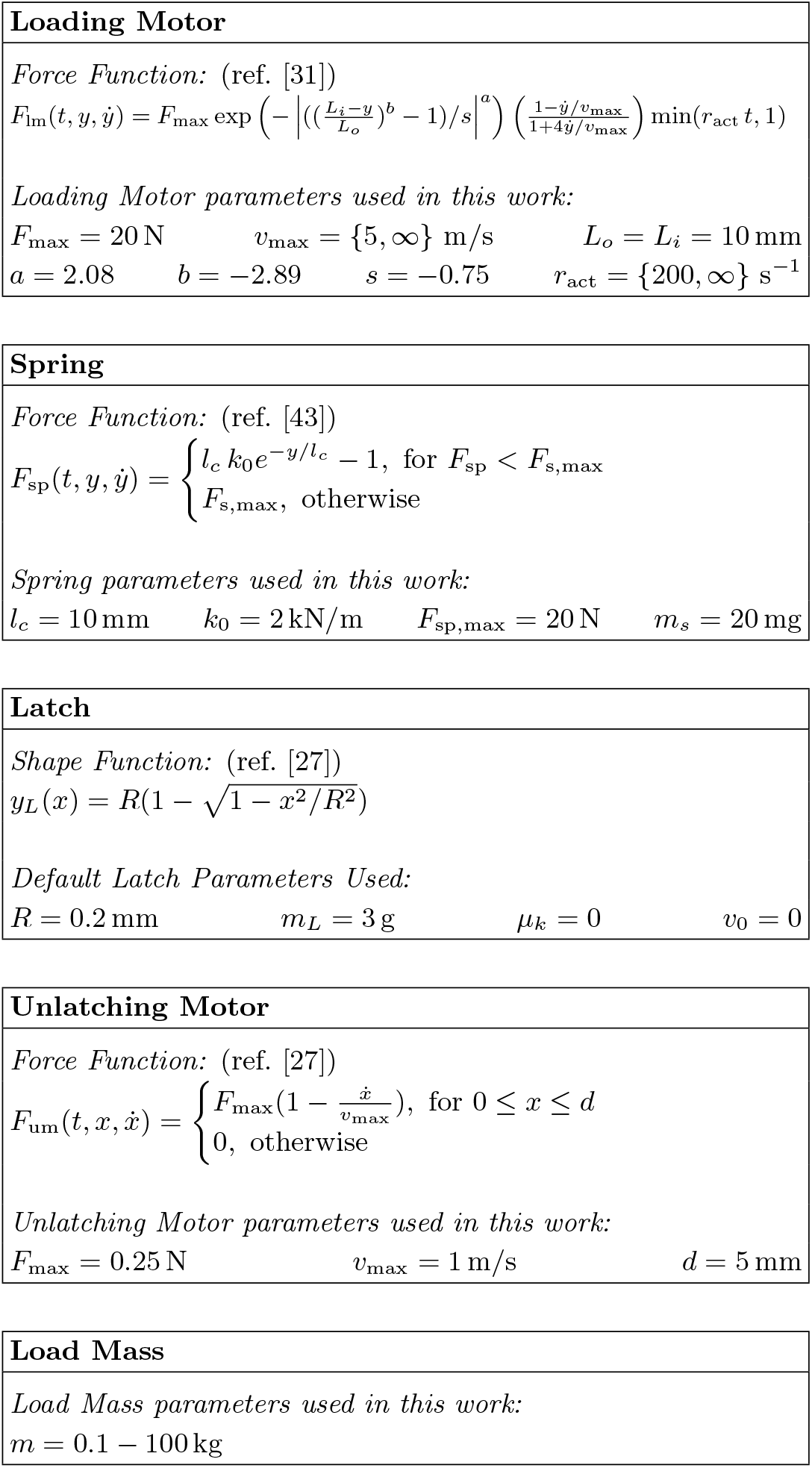
Mathematical description of the LaMSA components and default parameter values used in this work. The parameters were selected based on the range of characteristic forces, lengths, and velocities for biological LaMSA systems [27, 33].

|  |
| --- |
| **Loading Motor** |
| *Force Function:* (ref. [31])<br>$F_{\text{lm}}(t, y, \dot{y}) = F_{\text{max}} \exp\left(-\left\vert \left(\frac{L_i - y}{L_o}\right)^b - 1\right/s\right\vert ^a\right) \left(\frac{1 - \dot{y}/v_{\text{max}}}{1 + 4\dot{y}/v_{\text{max}}}\right) \min(r_{\text{act}} t, 1)$<br>*Loading Motor parameters used in this work:*<br>$F_{\text{max}} = 20 \text{ N} \quad v_{\text{max}} = \{5, \infty\} \text{ m/s} \quad L_o = L_i = 10 \text{ mm}$ $a = 2.08 \quad b = -2.89 \quad s = -0.75 \quad r_{\text{act}} = \{200, \infty\} \text{ s}^{-1}$ |
| **Spring** |
| *Force Function:* (ref. [43])<br>$F_{\text{sp}}(t, y, \dot{y}) = \begin{cases} l_c k_0 e^{-y/l_c} - 1, & \text{for } F_{\text{sp}} < F_{\text{s,max}} \\ F_{\text{s,max}}, & \text{otherwise} \end{cases}$<br>*Spring parameters used in this work:*<br>$l_c = 10 \text{ mm} \quad k_0 = 2 \text{ kN/m} \quad F_{\text{sp,max}} = 20 \text{ N} \quad m_s = 20 \text{ mg}$ |
| **Latch** |
| *Shape Function:* (ref. [27])<br>$y_L(x) = R(1 - \sqrt{1 - x^2/R^2})$<br>*Default Latch Parameters Used:*<br>$R = 0.2 \text{ mm} \quad m_L = 3 \text{ g} \quad \mu_k = 0 \quad v_0 = 0$ |
| **Unlatching Motor** |
| *Force Function:* (ref. [27])<br>$F_{\text{um}}(t, x, \dot{x}) = \begin{cases} F_{\text{max}}(1 - \frac{\dot{x}}{v_{\text{max}}}), & \text{for } 0 \leq x \leq d \\ 0, & \text{otherwise} \end{cases}$<br>*Unlatching Motor parameters used in this work:*<br>$F_{\text{max}} = 0.25 \text{ N} \quad v_{\text{max}} = 1 \text{ m/s} \quad d = 5 \text{ mm}$ |
| **Load Mass** |
| *Load Mass parameters used in this work:*<br>$m = 0.1 - 100 \text{ kg}$ |

### Appendix B: Derivation of the Model

Here we consider a simplified Latch Mediated Spring Actuated (LaMSA) system:

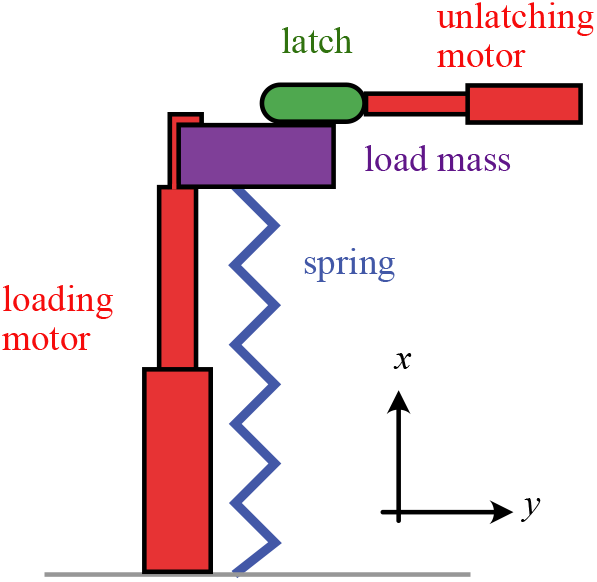

Our goal is to derive a single ordinary differential equation describing x(t) of the latch while it is in contact with the load mass.

#### Setting Up the Problem

Let us approximate the load mass and latch as point masses and draw isolation diagrams. In this model, we will consider the latch to have some shape that governs the unlatching process. Therefore, we have some function *y*_*L*_(*x*) that determines the curve of the latch.

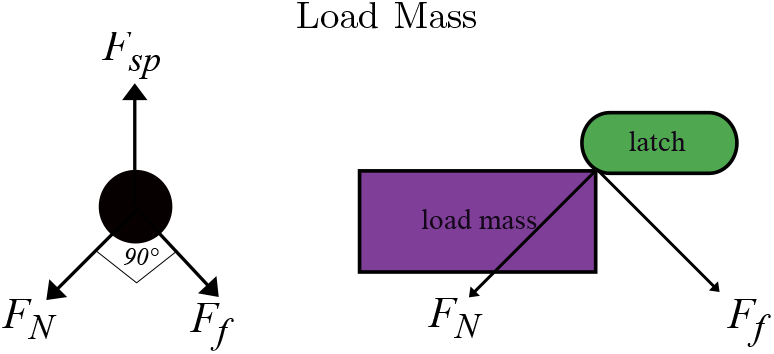

Variables:

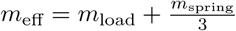: the effective mass of the load mass and spring mass combined

*F*_sp_ : force exerted by the spring on the load mass

*μ*_*k*_ : coefficient of friction between the load mass and the latch

*F*_*N*_ : normal force

*F*_*f*_ : force of friction between latch and load mass

*θ* : The angle between the normal force vector and the vertical

Among these variables, we’ll consider the following to be given: *F*_sp_, *μ*_*k*_

With names for our variables, we can write Newton’s second law to get the following:

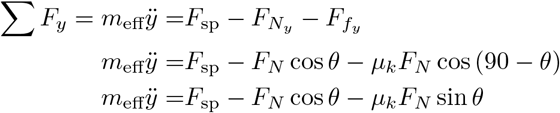

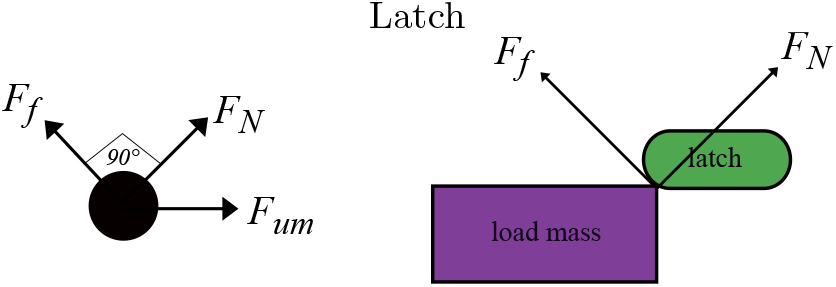

Variables:

*m*_*L*_ : mass of the latch

*F*_um_ : force of the unlatching motor pulling the latch away

*F*_*N*_ : Normal force from load mass on latch

*F*_*f*_ : Friction forcse from load mass on latch

*y*_*L*_(*x*) : function describing the latch geometry

Known Values:

*m*_*L*_ : mass of the latch

*F*_um_ : force of the unlatching motor pulling the latch away

*y*_*L*_(*x*) : function describing the latch geometry

We get the following equations from Newton’s 2nd Law:

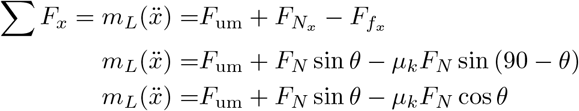

#### Rewriting Unknowns in Terms of Other Variables

We will use these replacements later in the derivation.

##### Rewriting *ÿ*

Our goal is to get a differential equation for 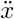, but we will end up with *ÿ* in our equations. So, we can use the following to rewrite *ÿ* in terms of 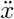 and the latch curve:

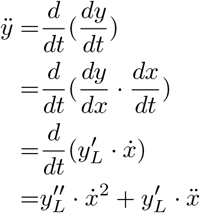

##### Rewriting tan *θ*

We will need to replace tan *θ* later in the derivation.

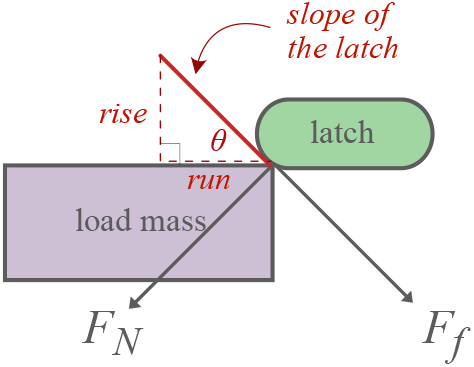

Because the latch geometry is described by the function *y*_*L*_, the slope of the latch is described by the derivative 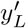.

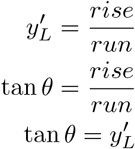

#### Solving for 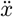

##### Solving for *F*_*N*_ in each equation

Recall that we obtained the following two equations from applying Newton’s 2nd Law to both :

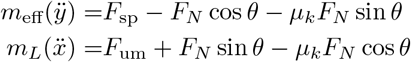

Let us replace *ÿ* with the expression we obtained in the previous section. Now we have:

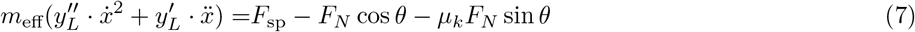

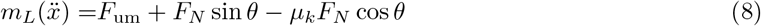

The only element that is not known is *F*_*N*_. Let us eliminate it by solving for *F*_*N*_ in both equations. Solving for *F*_*N*_ in Eq. (7) gives us:

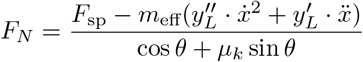

Solving for *F*_*N*_ in Eq. (8) gives us:

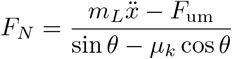

##### Expressing *F*_*N*_ without using 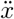

While it is our ultimate goal to solve for 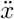, a side goal that is useful for determining the end of the unlatching phase is obtaining an expression for *F*_*N*_ that does not include 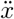.

We can achieve this by taking Eqs. (7) and (8) from the previous section, isolating 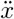 in each, and setting them equal to each other.

Rearranging Eq. (7):

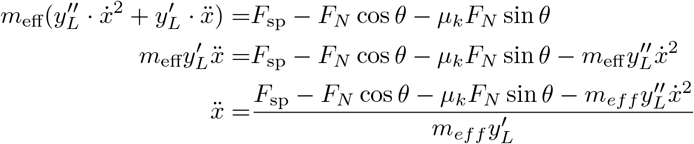

Rearranging Eq. (8):

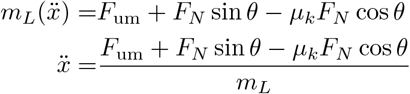

Setting these equal to each other, isolating *F*_*N*_ :

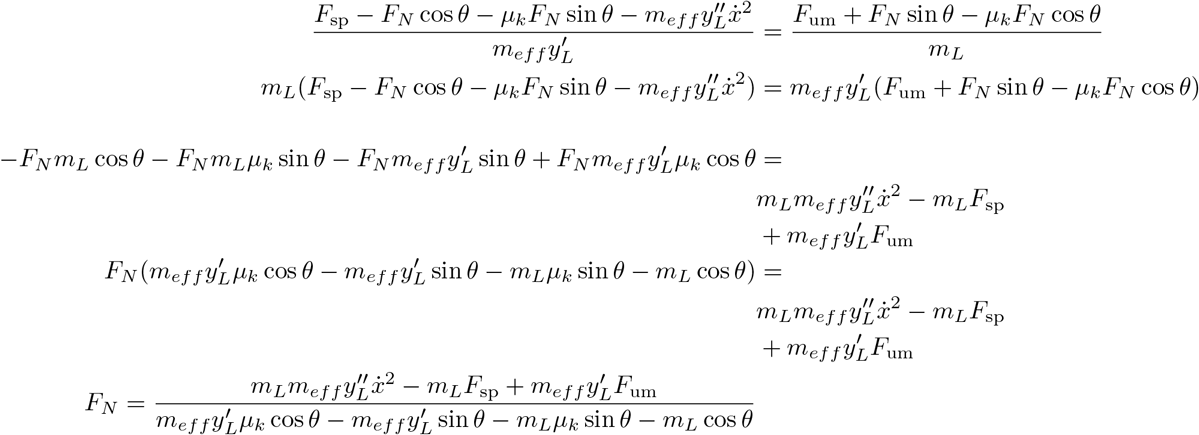

It’s somewhat inconvenient to have *θ* in this expression, so we can make the following substitutions based on the geometry of our problem:

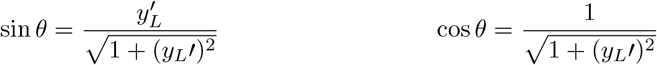

Plugging these in:

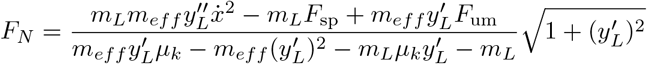

Great! Now that we have this, we’ll resume with our other goal, of solving for 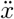.

##### Solving for 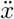

With two expressions for *F*_*N*_ in Eqs. (7) and (8), we can set them equal to each other to solve for 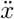:

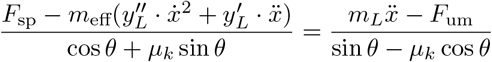

Cross-multiply to get:

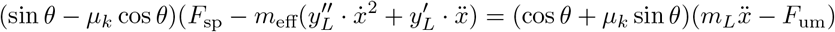

Expanding:

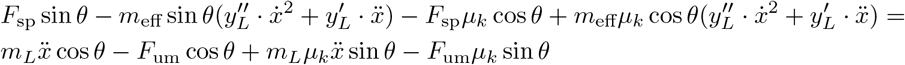

Divide both sides by cos *θ*:

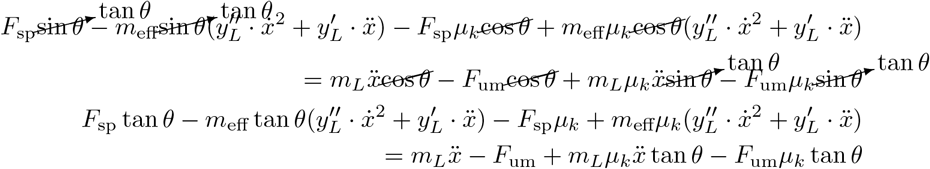

We can replace tan *θ* with 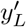:

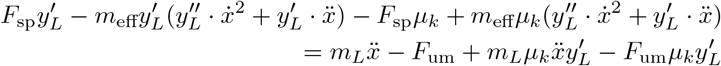

If we expand the equation, move all terms that contain 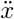 and 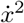 to one side, and factor out 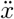 and 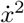, we get:

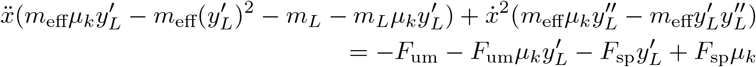

Now, let us solve for 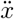 and regroup some terms to arrive at the final result:

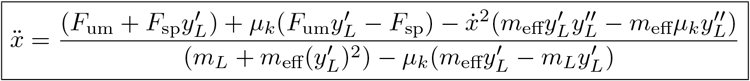

And if there is no friction such that *μ*_*k*_ = 0:

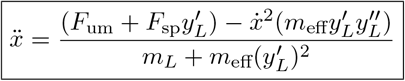

### Appendix C: Rotational Model

To model rotational motion, we can no longer assume *F*_*spring*_ = *m*_*proj*_ *· ÿ* as we do for the case of linear motion. Instead, there is a changing mechanical advantage as a function of the angle between the spring and the lever (pictured below) that complicates the dynamics of the system.

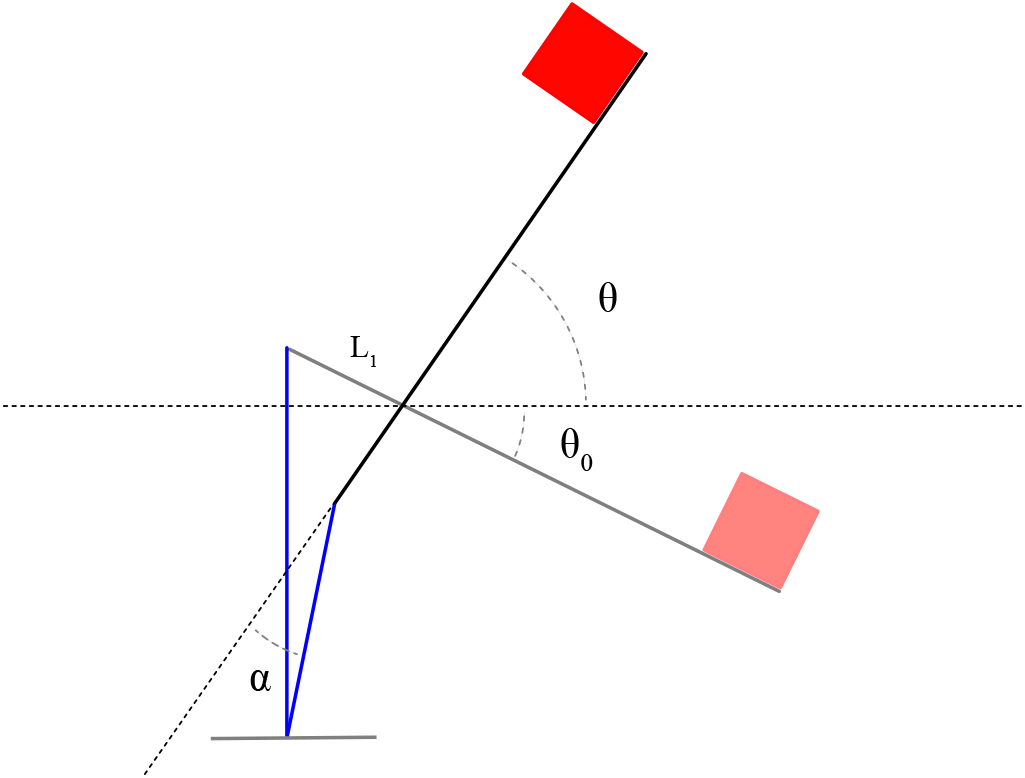

However, if we derive expressions for effective spring force and projectile mass *F*_eff_ and *m*_eff_ as functions of the displacement of the spring, we can use these expressions in our linear model to accurately describe rotational motion. In the following section, we derive these expressions.

We begin by writing Newton’s second law for rotational motion:

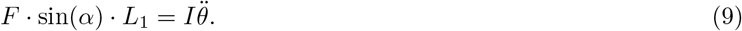

Our end goal is to write this as

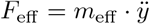

where *y* is the displacement of the spring.

First, we will write 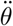 in terms of *ÿ*. There is a complex exact relationship between *y* and *θ* that an interested reader can calculate using the law of cosines a few times, but it is very well approximated by *y* = *L*_1_ *·* sin(*θ*). Using this, we find

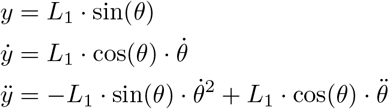

Rearranging these equations, we find

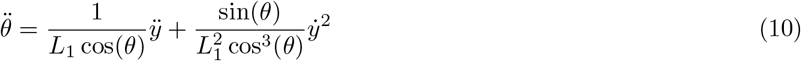

Now, if we substitute equation 10 into equation 9, we have

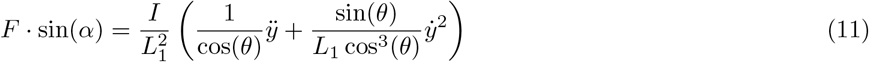

Next, we find 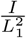. Using the parallel axis theorem, we find the moment of inertia of the lever (a rod with uniformly distributed mass *m*) and projectile (a point mass *M*) about the axis of rotation will be

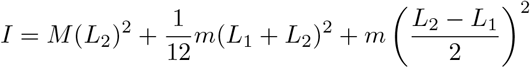

Dividing by 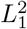 and substituting the effective mechanical advantage 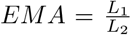, we have found an intermediate mass quantity *m*_int_ such that

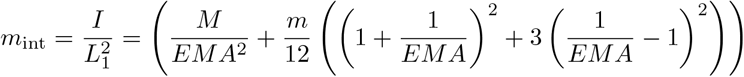

and we can rewrite equation 11 as

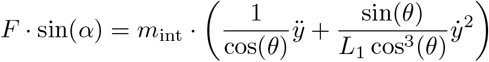

or

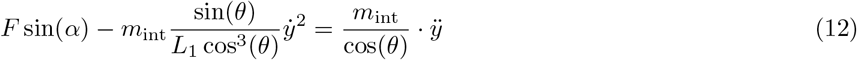

Equation 12 is in the desired form, so we can now extract

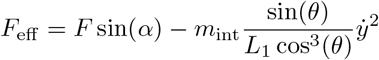

and

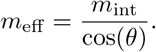

